# RNAseq sheds light on “who is doing what” in the coral *Porites lutea*

**DOI:** 10.1101/2025.01.09.632055

**Authors:** Kshitij Tandon, Juntong Hu, Francesco Ricci, Linda Louise Blackall, Mónica Medina, Michael Kühl, Heroen Verbruggen

**Affiliations:** School of BioSciences, The University of Melbourne, Melbourne, VIC 3010, Australia; Melbourne Integrative Genomics, The University of Melbourne, Melbourne, VIC 3010, Australia; Department of Microbiology and Immunology at the Peter Doherty Institute of Infection and Immunity, The University of Melbourne, Melbourne, VIC 3010, Australia; Department of Microbiology, Biomedicine Discovery Institute, Monash University, Clayton, VIC 3800, Australia; Securing Antarctica’s Environmental Future, Monash University, Clayton, VIC 3800, Australia; Department of Biology, Pennsylvania State University, University Park, PA 16802, USA; Marine Biological Section, Department of Biology, University of Copenhagen, DK-3000 Helsingør, Denmark; CIBIO, Centro de Investigação em Biodiversidade e Recursos Genéticos, InBIO Laboratório Associado, Campus de Vairão, Universidade do Porto, 4485-661 Vairão, Portugal

**Author notes:** Corresponding author: Kshitij Tandon.

## Abstract

Global decline of coral reefs due to climate change necessitates nature-based protection strategies for these crucial ecosystems. Developing such strategies requires a thorough understanding of the complex roles and interactions occurring within the coral holobiont. Using RNAseq, we investigated the active microbiome of healthy stony coral *Porites lutea*, focusing on the coral tissue, the green endolithic algal layer (*Ostreobiu*m layer), and the deeper coral skeleton. We identified distinct, metabolically active communities within these compartments and highlight substantial metabolic redundancy across carbon, nitrogen, and sulphur pathways. Our study provides first transcriptomic evidence of *Ostreobium’s* ability to transfer fixed carbon to other holobiont members and the coral host. Additionally, we highlight critical roles of diverse coral holobiont members in nutrient cycling and maintaining homeostasis through scavenging of reactive oxygen and nitrogen species. This study provides novel molecular-level understanding of the functional roles played by diverse coral holobiont members in their respective compartments and underscores that corals harbour distinct microbiomes with wide-ranging functions.

## Introduction

Coral reefs owe their success to the symbiosis between reef-building, stony corals (Scleractinia) (*1–3*) and a large diversity of closely associated bacteria, archaea, viruses, and microeukaryotes, collectively termed the coral holobiont (*3–6*). These symbionts *sensu lato* encompass organisms spanning the full spectrum of host-microbe interactions, from mutualistic partners that enhance coral fitness to opportunistic pathogens, as well as commensals and other microorganisms whose ecological roles may shift along this continuum depending on environmental conditions. While the most well-characterized symbiosis in corals involves photosynthetic, endosymbiotic microalgae in the family Symbiodiniaceae residing in the gastrodermal tissues of the animal host (*7–9*), a compendium of prokaryotes associated with corals has identified 39 bacterial and 2 archaeal phyla (*10*). This diverse microbial assemblage is not only distinct between different compartments within the coral (*11*), but fine-scale mapping of the coral skeleton has revealed a stratified community across its depth (*12*–*14*). In recent years, other microeukaryotes besides Symbiodiniaceae, including endolithic algae of the genus *Ostreobium*, fungi, and apicomplexans have also garnered attention (*15*, *16*).

The diversity and composition of coral holobiont members have been investigated through metabarcoding of ribosomal marker genes (16S rRNA, 18S rRNA, ITS2) (*17–19*) (*20*, *21*), providing invaluable insights into the diverse and structured symbiont composition *sensu lato* within coral compartments (*12*, *13*, *22*, *23*). Combined culture-based and molecular approaches have expanded our understanding of the functions of coral holobiont members in maintaining coral health and resilience (*24–28*). Symbiodiniaceae, in particular, play a crucial role by supplying corals with photosynthates, meeting a significant portion of their energy demand (*29*). Coral-associated bacteria and archaea have been implicated in a myriad of functions essential for sustaining coral health and resilience, including stress alleviation through scavenging of reactive oxygen and nitrogen species (ROS and RNS), protection against pathogens and nutrient cycling (*24–26*, *28*, *30–32*). Coral skeleton-dwelling endolithic green algae of the genus *Ostreobium,* which form conspicuous green bands beneath the coral tissue, have also been suggested to support corals by providing photosynthates during coral bleaching (*33–35*), but we lack a detailed understanding of the metabolic interaction between the coral animal and *Ostreobium* (*15*).

Most of our present knowledge on the role of microbiota for coral function and health comes from amplicon and metagenomic sequencing studies, with potential functions inferred using correlative, bioinformatic analyses. Despite these advances and the extensive information now available on the gene-encoded functional potential of microbiome and microeukaryotic members through genomic and metagenomic studies, a significant knowledge gap persists regarding actual microbial metabolic activity and its spatial organisation within the coral holobiont. Genomic data alone cannot predict the actual contributions of specific functions by symbionts *in situ*, as genes encoding these functions may be variably expressed or not expressed at all. RNAseq enables the analysis of the pool of actively expressed genes in an environmental sample, highlighting the ecophysiologically relevant processes occurring at the time of sampling (*36*). Therefore, taking a transcriptomics view can provide insights into which community members perform which functional roles.

Obtaining a molecular level understanding of the functional roles of holobiont members is crucial especially given the rapid decline of coral reefs worldwide due the adverse effects of climate change (*37*). Addressing this knowledge gap will also be beneficial for developing effective nature-based interventions to protect coral reefs, including improved assisted evolution strategies and development of coral probiotics (*2*, *38*). Both approaches rely on a comprehensive understanding of the roles played by diverse microbial symbionts in maintaining coral health.

In this study, we sought to answer the fundamental question of ’who is doing what’ in a healthy coral holobiont across different compartments. Using RNAseq, we identified the active symbiont community of the coral *Porites lutea*, characterizing their specific activities and locations within the coral, from the tissue to the endolithic algal layer (Ostreobium-layer) and deeper coral skeleton. We focused on key nutrient cycling pathways, including carbon fixation (C), photosynthesis (P), nitrogen (N) and sulfur (S) metabolism, sugar export, and the scavenging of reactive oxygen and nitrogen species (ROS and RNS).

## Methods

### Coral sampling and colony fragmentation

Fragments from six healthy-looking, *Porites lutea* colonies, spaced at least 15 m apart, were collected at low tide (1-2 m depth) from the research zone of the Heron Island reef flat, southern Great Barrier Reef (23°44′S, 151°91′E), in October 2022. Four fragments (from colonies we numbered 1, 3, 4 and 5) were collected during daytime, while two (colonies 6 and 7) were collected five-six hours after sunset. All collected colonies exhibited a visually discernible *Ostreobium-layer* in the coral skeleton . 10-50 mm below the coral tissue layer (Fig. 1A). The fragments were collected using a sterile hammer and chisel and were placed in separate sterile zip-loc polyethylene bags in seawater and flash frozen in liquid nitrogen immediately upon collection. A 5-mm thick slice was cut from the frozen samples, perpendicular to the vertical growth axis, using a diamond saw with a continuous flow of sterile filtered (0.2 µm) seawater. Later, pliers were used to obtain fragments from three distinct layers, i.e., the coral tissue, the green *Ostreobium*-layer, and the tan fraction of the skeleton below the green band (termed the skeleton layer). During this process, the material was kept frozen by repeated immersion in liquid nitrogen. Each layer was kept frozen at -80°C, transported to the University of Melbourne on dry ice, and processed immediately upon arrival.

**Fig. 1.**
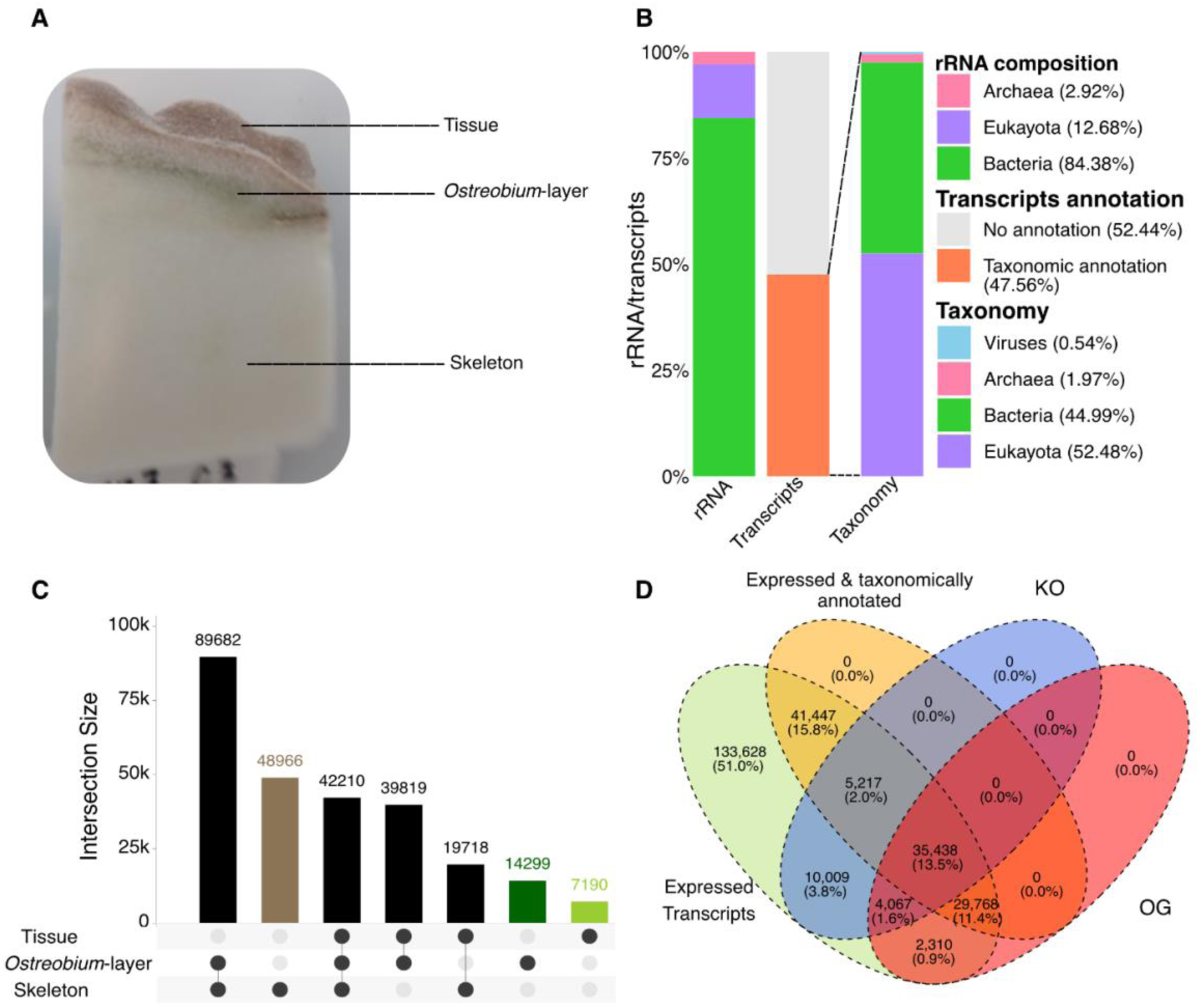
Taxonomic and functional overview based on rRNA and transcriptome data. **A)** illustration of a coral fragment depicting the three compartments: Tissue, Ostreobium-layer and Skeleton. B) Bar plots showing overall community composition based on rRNA and transcriptome data. C) UpSet plot illustrating shared and unique transcripts across different coral compartments. D) Venn diagram showing counts and percentage of transcripts which are deemed expressed, expressed and taxonomically annotated, annotated at KO and at OG levels.

### Total RNA extraction

For RNA extraction, samples of individual layers were ground to a fine powder in liquid nitrogen using pre-cooled mortars and pestles. Approximately 200 mg of powdered tissue samples and between 300-500 mg of powdered *Ostreobium-layer* and skeleton layer were used for RNA extraction with 1 mL of pre-cooled TRIzol reagent (Invitrogen, USA). Total RNA was extracted following the manufacturer’s protocol with some modifications (**see supplementary file 1 for details**). Quantification of total RNA was conducted using the Qubit RNA HS Assay kit (Thermo Fisher Scientific, USA).

### RNA sequencing and data-preprocessing

We did not perform any ribosomal RNA depletion due to the dual aim of sequencing mRNA and rRNA from both eukaryotes and prokaryotes simultaneously and profiling the transcriptionally active microbial and microeukaryotic community composition using 16S and 18S rRNA abundance (see below). Total RNA libraries were prepared for 36 samples (6 colonies x 3 layers x 2 replicates per layer) and sequenced on the Illumina NovaSeq platform by Azenta (Suzhou, China). Sequencing yielded approximately 23 million reads per sample with 2x150 bp reads.

Raw reads were quality checked and trimmed using FastQC v0.11.9 (https://www.bioinformatics.babraham.ac.uk/projects/fastqc) and Trimmomatic v0.39(*39*), respectively. Trimmomatic parameters were as follows: *TruSeq3-PE.fa:2:30:5 HEADCROP:5 SLIDINGWINDOW:4:20 MINLEN:50*. Quality-filtered reads were first used to identify and filter out rRNA reads using SortMeRNA v4.3.6(*40*) with Silva v138.1 LSU and SSU (*SILVA138.1 LSU and SSU NR99 database, last accessed Feb 10, 202*3)(*41*). Reads identified as rRNA were taxonomically classified using PhyloFlash v3.4(*42*) (with *almost-everything*), which rapidly reconstructs the SSU rRNAs (NTUs, nearest taxonomic units) and taxonomically annotates them using Silva 138.1 NR99 SSU database. Reads identified as rRNA were removed from the raw-read set, and non-rRNA reads were mapped to the *Porites lutea* genome (accessed from www.reefgenomics.org) using bowtie2 v2.3.4.1(*43*) and samtools 1.17(*44*) to identify and remove host-related reads.

High-quality, non-rRNA and non-host reads were co-assembled using Trinity v2.15.1(*45*) with parameters *--min_contig_length 250 and --no_salmon*. Taxonomic annotation of assembled transcripts was obtained using the lowest common ancestor approach by performing *blastx* (*46*) searches against the NR database (last accessed April 2023) with diamond v2.1.6.160(*47*), using an *e-value cutoff of 1e-5* and processed through MEGAN6 community edition v6.24.20, using the *megan-map-Feb2022-ue.db* and meganising the *daa* file using *daa2info* (*48*). This process was used to identify and remove any transcripts annotated as Scleractinia and other Eumetazoa. The remaining transcripts were quantified using RSEM (RNA-Seq by Expectation and Maximisation)(*49*), implemented in Trinity through the utility script *align_and_estimate_abundance*.*pl*. This approach generated both gene and isoform expression profiles. Isoform-level expression data was used for the analysis. Expression values per transcript were averaged across replicates for each layer (tissue, *Ostreobium*-layer and skeleton).

### Community composition analysis

Taxonomic classification of NTUs assembled from rRNA reads was conducted to profile the abundance of transcriptionally active community members. NTUs classified as “Chloroplast” or “Mitochondria” were removed. Abundance values were converted to relative abundance for compositional analysis using barplots and log-transformed for diversity analysis using PCoA. These analyses were performed using R packages phyloseq v1.42.0(*50*), metagMisc v0.5.0 (https://github.com/vmikk/metagMisc) and ggplot2 v34.3 (*51*). We used the quantified and taxonomically annotated transcripts assembled from mRNA reads to profile the community composition encoding for different functional guilds based on the transcriptome data and compared them to rRNA based analysis using the same approach.

### Functional analysis

To ascertain functional roles, we first filtered transcripts using Trimmed Mean of the M (TMM) values of 0.1 and presence in at least 2 samples to identify truly expressed transcripts. This approach filters out i) any transcripts from potential residual DNA or very low expressed transcripts, ii) transcripts originating from spurious mapping of reads. Transcripts deemed truly expressed were then grouped in a compartment-specific manner to obtain tissue, *Ostreobium*-layer and skeleton-layer specific transcriptome. These compartment specific transcripts were processed through Trinotate v4.0.0(*52*) and Transdecoder v4 (https://github.com/TransDecoder/TransDecoder) to identify the longest open reading frames and predict the proteins from transcripts ≥50 amino acids. Predicted proteins were functionally annotated using a combination of EggNOG mapper v2.1.12(*53*) with database eggnogDB v5.0.2(*54*) with *e-value cutoff 1e-5*, ghostkoala (*55*) with database *c_family_euk+genus_prok+viruses* parameters and Interproscan v5.63-95.0(*56*) with *e value cutoff 1e-5* to obtain a comprehensive understanding of functional orthologous groups (EggNOG and COG), KEGG pathways, including Kegg Ontology, KOs and protein families (pfam). Metabolic markers for carbon-fixation (C), photosynthesis (P), nitrogen (N) and sulfur (S) metabolism were identified from a previous study (*57*). Additionally, we focussed on genes involved in sugar export and scavenging of ROS and RNS. Expression profiles of transcripts functionally annotated with marker KOs or pfams were plotted using heatmaps using ComplexHeatmap(*58*) in R.

## Results and Discussion

### Corals harbor compartment-specific active symbiont community

One of our primary objectives was to profile the transcriptionally active community within the coral holobiont and compare these findings with the community composition obtained from rRNA analysis. To achieve this, we examined the total and active community composition across the coral tissue, the endolithic *Ostreobium-layer*, and the underlying coral skeleton, and compared it with their rRNA based community composition (**Fig 1A)**. For total community composition, we taxonomically analysed 28,717 rRNA NTUs (6,010 from tissue layer, 15,942 from *Ostreobium* layer, 26,712 from skeleton layer) belonging to Archaea (2.92% - 840 sequences), Eukaryota (12.68% - 3,644 sequences), and Bacteria (84.38% - 24,233 sequences) (**Fig 1B**). This observation offers essential insights into the diversity and composition of the coral holobiont. Unlike traditional amplification-based metabarcoding analyses, which are often biased (*59*) and fail to provide accurate quantitative distinctions between prokaryotes and eukaryotes, using rRNAs assembled from total RNA delivers a more precise and comprehensive overview of the community composition. Eukaryotes (Amorphea, SAR, Archaeplastida) were the dominant groups in the coral tissue (**Supplementary Fig S1A**) and the *Ostreobium layer* (**Supplementary Fig S1B**), while the deeper skeleton was dominated by Bacteria (Proteobacteria, Planctomyceota, Desulfobacteria, and Cyanobacteria) (**Supplementary Fig S1C**). Our study shows that bacterial diversity is highest in the skeleton, where the majority of bacterial NTU sequences were assembled, confirming previous reports of greater bacterial richness in the skeleton compared to tissue compartments (*13*, *19*).

After initially evaluating NTUs based on rRNAs, we extended our analysis to the remaining RNA sequence annotations. To profile active microbial community composition, we used a curated set of 371,193 taxonomically annotated transcripts, selected from a total of 780,431 assembled sequences. After excluding 111,367 transcripts belonging to Scleractinia, Chordata, and Eumetazoa, 259,826 (47.56%) transcripts remained. These transcripts represent Eukaryotes (52.48% - 136,366), Bacteria (44.99% - 116,900), Archaea (1.97% - 5,131), and viruses (0.54% - 1,429) (**Fig 1B**) and revealed a high relative abundance of alveolates, the eukaryotic group containing Apicomplexa and dinoflagellates, including Symbiodiniaceae in coral tissue (**Supplementary Fig S2A)**. Green algae, Streptophyta and Chlorophyta, dominated the *Ostreobium layer* and deeper skeleton. (**Supplementary Fig S2B and C**). The high relative abundance of dinoflagellate (Symbiodiniaceae) expressed transcripts in coral tissue is unsurprising, as they are the most dominant microeukaryotes in the coral tissue, meeting most of the coral’s energy demands (*60*). However, we identified dinoflagellate and Apicomplexa transcripts in all compartments. Apicomplexans have been found in healthy coral colonies across the geographic distribution and taxonomic diversity of corals, but their role is yet to be characterized (*16*, *61*). Bacteria and Fungi were present in all compartments, with bacteria as dominant members of the skeletal microbiome (**Supplementary Figure S2C)**. Proteobacteria, Cyanobacteria and Actinobacteria were the dominant bacterial phyla in all coral compartments. Members of the phylum Proteobacteria are some of the most widespread coral microbiome members (*10*, *17*, *62*). Cyanobacteria have also previously been identified as the dominant prokaryote group in the *Ostreobium layer* and deeper skeleton of *P. lutea* from the Red Sea (*63*). Obtaining annotation for less than 50% of the total assembled transcripts highlights the immense diversity of uncharacterised symbionts associated with *P. lutea,* despite it being one of the best investigated coral species with several metagenomic and physiological studies (*24*, *25*, *64*, *65*). These findings align with earlier studies using the metatranscriptomics approach to profile transcriptionally active microbiomes in other ecologically diverse systems, including soil (*66–68*), sponge (*69*) and ocean microbiomes (*57*, *70*).

### Functional annotation of transcripts remains a bottleneck

Analysis of active metabolic pathways in a healthy coral holobiont revealed 261,884 truly expressed transcripts through TMM filtering. These transcripts were distributed across the three investigated compartments, with the highest overlap observed between the *Ostreobium layer* and deeper skeleton (89,682 shared transcripts). Additionally, a substantial number of uniquely expressed transcripts were identified in the deeper skeleton-layer (48,966), while the coral tissue exhibited the lowest count of uniquely expressed transcripts (**Fig. 1C**). Recent metabarcoding and metagenomic studies have highlighted the skeleton as the most diverse compartment of corals (*13*, *19*, *63*). In line with these observations, our coral skeleton samples revealed 178,580 uniquely predicted proteins from the expressed transcripts. However, only 30.64% (54,731) could be annotated using KO ids and 29.25% (52,252) using OGs (**Fig 1D**). The extraordinary uniqueness of skeletal proteins underscores the added value of sampling genomically underexplored environments, such as the coral skeleton. The low functional annotation observed in our dataset is consistent with a recent study on gene expression changes and community turnover of global oceans, which similarly found less than 25% of the transcripts assigned to KOs (*57*). Annotations based on COG revealed a notable proportion of proteins encoded by transcripts involved in signal transduction, intracellular trafficking, secretion, vesicular transport, post-translational modification, protein turnover, and chaperones (**Supplementary Figure S3)**. While most transcripts remained unannotated, the presence of these functional categories suggests potential microbial involvement in cellular communication and protein homeostasis within the coral holobiont. Although our data do not directly demonstrate nutrient or metabolite exchange, these functional signals are consistent with prior hypotheses that such microbial activities may support coral resilience, particularly in oligotrophic environments. Further targeted studies, including isotopic tracing or compartment-specific transcriptomics, will be needed to directly test these functional roles.

### Distinct photosymbionts drive carbon fixation in coral compartments

To reveal the functional roles of transcriptionally active holobiont members in healthy *P. lutea* coral, we identified key microbial processes, such as photosynthesis, carbon fixation, sugar export, and nitrogen and sulphur cycling. Our analysis of pathway-specific metabolic markers highlighted the taxonomic affiliations of transcripts and their expression profiles, shedding light on the active contributions of each microbial group to these essential processes. Among the most highly expressed genes were those associated with photosynthetic carbon fixation, including photosystem I (PsaA, 59 transcripts), photosystem II (PsbA, 67 transcripts) and the Calvin–Benson cycle enzyme, RuBisCO subunits (ribulose-1,5-bisphosphate carboxylase/oxygenase, *rbc*L (79) and *rbc*S (18) genes) (**Fig 2**). While rbcL can be expressed by chemotrophs (*71*), our taxonomic annotations did not identify chemotrophic affiliations, suggesting that carbon fixation in this context is predominantly photoautotrophic. These genes were actively transcribed by Symbiodiniaceae, stramenopiles, Bryopsidales (including the endolithic green alga *Ostreobium*), Cyanobacteria, Gammaproteobacteria, and unclassified Viridiplantae and Chlorophyta in different coral compartments. Symbiodiniaceae dominated expression in the tissue, Cyanobacteria were active throughout all compartments, and *Ostreobium* and other Ulvophyceae were the main contributors in the deeper skeleton layers (**Supplementary data file 2**). The co-expression of *rbc*L/*rbc*S with *psb*A and *psa*A supports the role of chlorophototrophy as the dominant carbon fixation strategy in the coral holobiont (*72–74*).

**Fig. 2.**
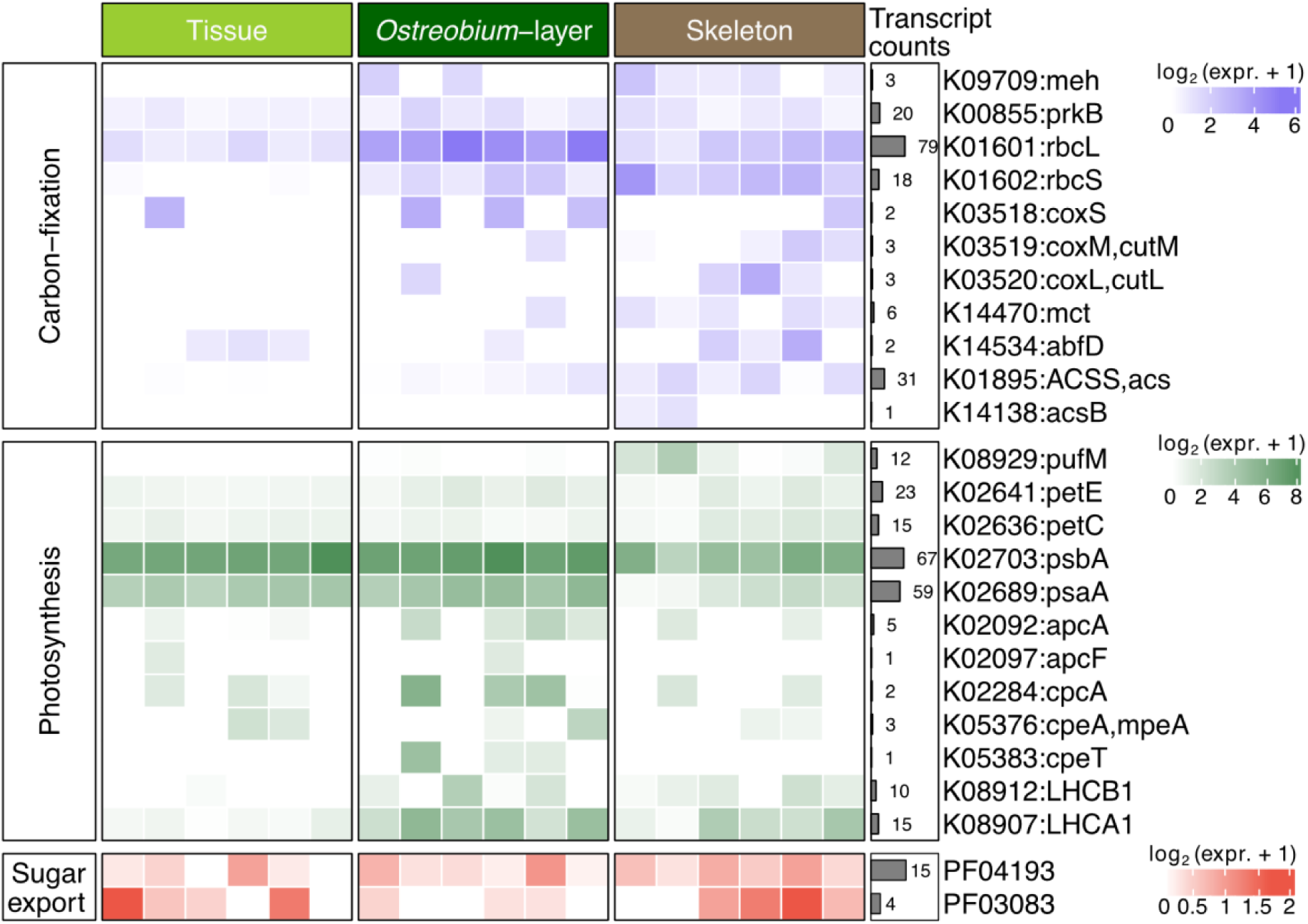
Carbon Fixation, Photosynthesis and Sugar Export Across Coral Compartments. Heatmap displaying the expression of marker genes for carbon fixation photosynthesis, and SWEET protein-mediated sugar export across various coral compartments. Expression values for both were averaged for the count of transcripts and log2(expression + 1) transformed.

In the skeleton, we also detected transcripts indicative of alternative carbon fixation pathways. The expression of *cox*L, *cox*M, and *cox*S, affiliated with Cyanobacteria and Proteobacteria, suggests potential activity of carbon monoxide dehydrogenase (CODH), enabling CO oxidation and energy generation through chemosynthesis. Additionally, transcripts for *acs* and *acs*B, encoding the acetyl-CoA synthase complex of the Wood–Ljungdahl pathway, were expressed in the skeleton and affiliated with anaerobic bacteria. This pathway enables fixation of CO_2_ and CO into acetyl-CoA, a central metabolite in microbial energy and biosynthetic pathways (*63*, *75*). These transcriptional signals reflect the metabolic versatility of skeleton-associated microbial communities and suggest adaptation to low-oxygen, possibly CO-rich microenvironments (*14*, *76*). Although we did not directly measure substrate availability or fluxes, the coexistence of phototrophic and anaerobic carbon fixation pathways implies that microbial communities play diverse roles in internal carbon cycling. These findings highlight the functional potential of coral-associated microbes to persist in spatially heterogeneous, nutrient-poor reef environments; however, further experimental work is needed to determine the direct impacts of these processes on coral physiology or holobiont productivity.

Besides *psa*A and *psb*A, we detected expression of several other photosynthetic marker genes across all three coral holobiont compartments. These included, *petC*, which encodes the Rieske iron–sulfur protein of the cytochrome b6f complex that links Photosystem II to Photosystem I, and *petE* ncoding plastocyanin, which facilitates electron transfer between the cytochrome b6f complex and Photosystem I. The expression of these genes supports the widespread presence of oxygenic phototrophs throughout the coral tissue, *Ostreobium* layer, and skeleton. transcripts for cyanobacteria-specific antenna proteins—including *apcA, apcF, cpcA, cpeA* and *cpeT*—were most abundant in the Ostreobium layer and deeper skeleton, suggesting a greater prevalence and transcriptional activity of Cyanobacteria in these compartments compared to the tissue (**Fig 2**). This pattern is consistent with taxonomic assignments based on both rRNA (**Supplementary Fig S1**) and transcriptome levels (Supplementary Fig S2) and aligns with previous findings showing an enrichment of cyanobacterial and eukaryotic phototrophs in the skeletal layers of *P. lutea* These antenna and photosynthetic genes, many of which are involved in the Calvin–Benson cycle, collectively accounted for more than 50% of the total carbon fixation gene transcripts in the skeleton. We also observed expression of *pufM*, a marker gene for anoxygenic photosynthesis that encodes a core component of the bacterial photosynthetic reaction centre. Expression of *pufM* was confined to the skeleton, suggesting the presence of obligate anaerobes, such as green sulfur and purple non-sulfur bacteria. This observation is in agreement with hyperspectral pigment imaging, which revealed pigment signatures consistent with anoxygenic phototrophs in the skeletal layers of *P. lutea* (*14*).

The observed compartment-specific expression patterns of photosynthetic genes reflect microbial adaptations to distinct light regimes within the coral holobiont and provide insight into the metabolic plasticity of associated phototrophs. The elevated expression of cyanobacterial antenna protein genes (*apcA*, *apcF*, *cpcA*, *cpeA*, *cpeT*) in the Ostreobium layer and deeper skeleton suggests that Cyanobacteria inhabiting these low-light regions may utilize alternative chlorophyll pigments, such as chlorophyll *d* and *f*, to absorb far-red light that penetrates coral tissue. This strategy likely enhances photosynthetic efficiency under reduced irradiance and aligns with previous reports of far-red light utilization by endolithic Cyanobacteria and *Ostreobium* (*77–79*).

Within the same skeletal microhabitat, the green alga Ostreobium appears to employ complementary adaptations, including the use of chlorophyll b and an expanded suite of light-harvesting complex (LHC) proteins in both photosystems (*15*). Genomic features such as Lhca1 gene duplication and the loss of photoprotective genes further reflect specialization to persist under light-limited conditions (80). In contrast, the tissue compartment, characterized by higher light availability, showed dominant transcriptional activity from Symbiodiniaceae and other phototrophs known to rely on chlorophyll *a*and *c*, consistent with adaptation to higher irradiance (81). The distinct expression of photosynthetic genes and pigment-related functions across coral compartments emphasizes the niche differentiation and photoadaptive strategies among coral-associated phototrophs. These spatial patterns of gene expression illustrate the functional diversity of the holobiont and suggest mechanisms by which microbial communities could contribute to energy capture across microhabitats with steep light gradients.

### Not only Symbiodiniaceae provide fixed carbon to the coral holobiont

To identify members of the coral holobiont involved in the export of photosynthetically fixed carbon compounds (photosynthates), we focused on genes encoding SWEET (“Sugars Will Eventually be Exported Transporter”) proteins. SWEETs are bidirectional sugar uniporters that facilitate the diffusion of sugars such as glucose along concentration gradients across biological membranes (*82*, *83* We identified transcripts encoding SWEET proteins, specifically associated with Pfam domains PF04193 and PF03083, across all three major compartments of the coral holobiont (**Fig 2**). Taxonomic annotation of these transcripts revealed that sugar exporters were predominantly eukaryotic, including Symbiodiniaceae, the endolithic alga *Ostreobium*, and unclassified green algae within the Chlorophyta (**Supplementary data file 2**). These organisms share a common capacity for oxygenic photosynthesis via the Calvin cycle, in which glyceraldehyde-3-phosphate (G3P), serves as a precursor for sugar biosynthesis, including glucose. Glucose is likely the principal carbohydrate exported via SWEET proteins (*84*), suggesting that these eukaryotic phototrophs contribute substantially to the pool of photosynthates available within the coral holobiont.

Interestingly, despite previous studies reporting sugar transporters in coral-associated bacteria (*26*), we did not detect transcripts encoding semiSWEET proteins, the prokaryotic homologs of SWEETs. This absence may reflect a biological partitioning of function; wherein eukaryotic symbionts serve as primary carbohydrate exporters. Alternatively, it could be a result of detection limits or the divergence of bacterial sugar transporters from characterized semiSWEETs. Regardless, our findings highlight a potentially central role for eukaryotes in sugar-based metabolic exchange among holobiont members.

Notably, we identified SWEET gene expression in *Ostreobium*, providing the first genomic and transcriptomic evidence that this skeleton-dwelling alga has the capacity to export sugars. These sugars may diffuse into the coral skeleton, potentially serving as an energy source for both the coral host and other microbial symbionts. This finding provides molecular support for prior radiolabeling studies using 14C and 13C-labeled photoassimilates, which demonstrated that fixed carbon from endolithic algae can be translocated into coral host lipids under both healthy and bleached conditions (*34*, *35*). The ability of *Ostreobium* to contribute sugars under thermal stress, when Symbiodiniaceae populations are diminished, suggests a compensatory role in mitigating coral energy deficits (*85*). This metabolic capacity may be especially important during bleaching events, offering a survival advantage by partially offsetting the loss of primary photosynthate sources.

In addition to phototrophs, we also detected the expression of SWEET-encoding transcripts in coral-associated fungi. This is the first direct molecular evidence of fungal participation in sugar transport within the coral holobiont. While the functional implications remain to be fully resolved, fungi are increasingly recognized for their roles in shaping coral-associated microbiomes and reef-scale biogeochemical cycling through nutrient processing and chemical interactions (*86*). Their involvement in fixed carbon exchange may reflect metabolic cross-feeding, carbon recycling, or interactions with other holobiont members, broadening our understanding of fungal contributions to holobiont function.

The identification of multiple eukaryotic lineages capable of translocating sugars underscores the complex and cooperative nature of carbon cycling within the coral holobiont. These sugar-exporting symbionts likely play critical roles in sustaining coral energy homeostasis. However, this capability may also pose risks: elevated extracellular sugar concentrations could unintentionally fuel the proliferation of opportunistic or pathogenic bacteria, potentially destabilize the microbial community and compromise coral health. Thus, while eukaryotic sugar exporters contribute to coral resilience, their activity may also influence disease susceptibility, highlighting the need for further investigation into the ecological and evolutionary consequences of carbon transfer dynamics in the coral holobiont.

### Skeleton dwelling microbes mediate nitrogen metabolism

Our analysis revealed active nitrogen metabolism within the skeleton of *P. lutea*, highlighting the role of skeleton-associated microbes in sustaining nitrogen cycling within the coral holobiont. Expression of canonical nitrogen fixation marker genes nifK, nifD and nifH was detected in both the *Ostreobium* layer and the deeper skeleton, suggesting active nitrogen fixation in these predominantly anoxic microenvironments (**Fig. 3A**). Nitrogen cycling is essential in coral reef ecosystems, where nutrient scarcity demands tight microbial recycling to support coral growth and productivity (*31*). While nitrogen-cycling microbes are widely documented in coral-associated microbiomes (*31*, *87*), our study identified Cyanobacteria as the predominant taxonomically resolved nitrogen fixers in *P. lutea*. This contrasts with earlier reports emphasizing heterotrophic nitrogen-fixing bacteria (*88–90*) and instead aligns with recent genome-resolved metagenomic studies showing low diversity of nitrogen-fixing taxa in this species (*24*, *25*). It is important to note that a substantial proportion of transcripts remain unclassified, and additional nitrogen-fixing taxa may be present but undetected. Nonetheless, the predominance of cyanobacterial transcripts underscores their likely importance in supporting nitrogen-limited growth of Symbiodiniaceae (*9*, *91*), and highlights their key role in the coral holobiont’s nitrogen economy.

**Fig 3.**
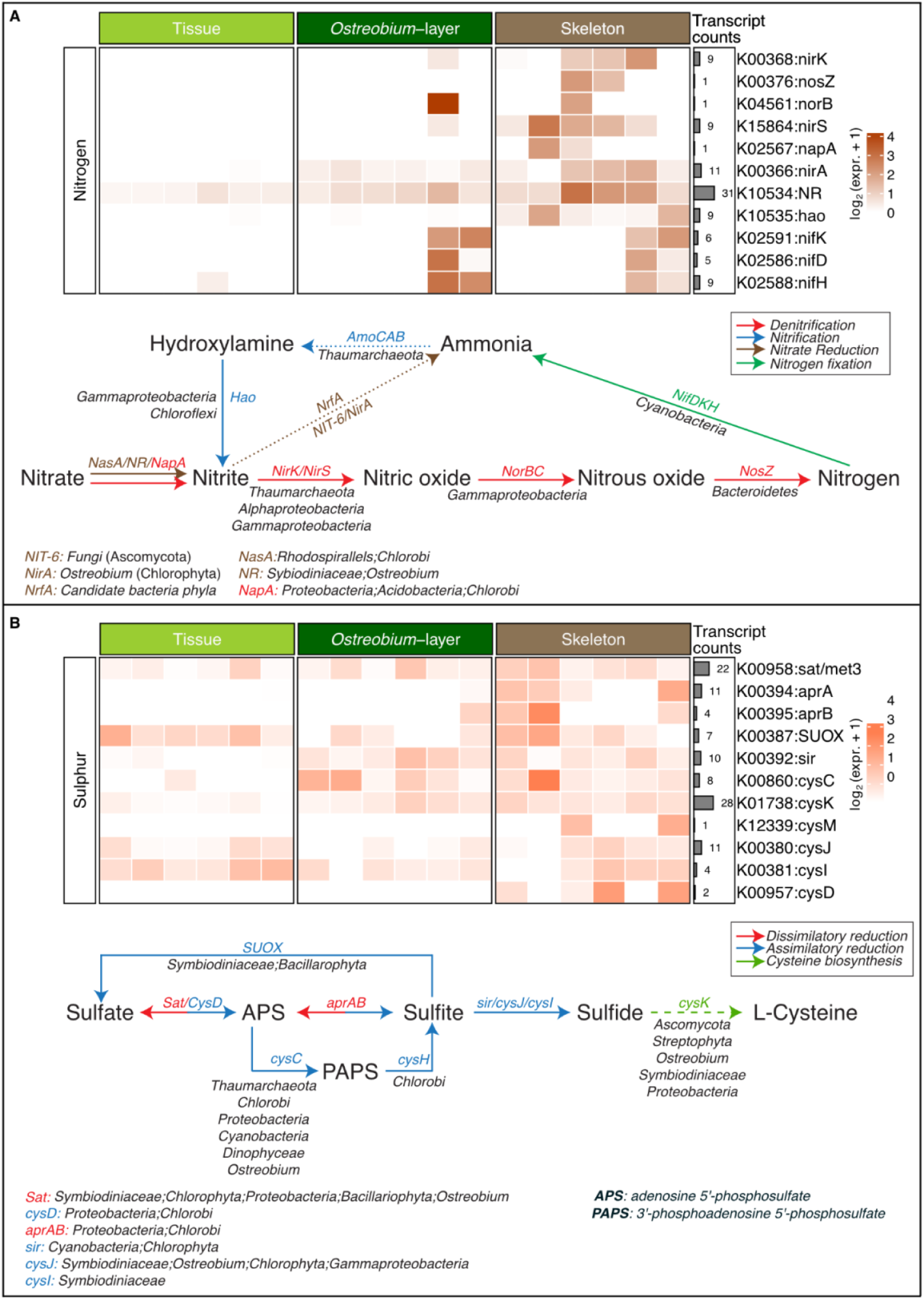
Transcriptome-based expression profiles of Nitrogen and Sulfur metabolism across coral compartments. A) Heatmap showing the expression profiles of marker genes involved in nitrogen metabolism across coral compartments, along with a bar plot indicating the count of transcripts. The nitrogen cycle is illustrated, highlighting subpathways and contributing community members. B) Heatmap showing the expression profiles of marker genes involved in sulphur metabolism across coral compartments, along with a bar plot indicating the count of transcripts. The sulphur metabolism pathway is illustrated, highlighting subpathways and contributing community members. Expression values for both were averaged and log2(expression + 1) transformed.

We also identified nitrification, the aerobic microbial process converting ammonia to hydroxylamine and subsequently to nitrite and nitrate, to be ubiquitous across all coral compartments (**Fig 3A**). This process is vital for nitrogen turnover and ammonium regulation, reducing toxicity, and maintaining nitrogen turnover within the holobiont, with implications for both coral host and symbiont physiology (*31*). Transcripts encoding nitrification-related genes were primarily assigned to Thaumarchaeota, Gammaproteobacteria, and Chloroflexi, particularly in the deeper skeleton (**Fig 3B and Fig 5**). These taxa likely contribute to ammonia oxidation, although unclassified transcripts and incomplete database coverage suggest that additional nitrifying microbes may be involved.

In contrast, denitrification was spatially restricted to the *Ostreobium* layer and deeper skeleton compartments, consistent with oxygen-limited conditions in these compartments. We detected expression of key denitrification marker genes (*napA, nirS, norB, and nosZ*) in both eukaryotic phototrophs (Symbiodiniaceae, Ostreobium) and bacterial symbionts, including Rhodospirillales, Chlorobi, Proteobacteria, and Acidobacteria (**Fig. 3A**; **supplementary data file 3**). The subsequent steps in denitrification, conversion of nitrite to nitric oxide, nitrous oxide, and nitrogen during denitrification were most commonly attributed to Thaumarchaeota, and Proteobacteria (α- and γ-), and Bacteroidota. Importantly, we also detected fungal (Ascomycota) transcripts encoding NIT-6 gene, involved in dissimilatory nitrate reduction to ammonium (DNRA), providing evidence for fungal contributions to nitrogen recycling within the holobiont.

The interplay between nitrification, denitrification, and DNRA, particularly within the skeleton, appears crucial for maintaining nitrogen homeostasis. DNRA may dominate under low-oxygen conditions (e.g., at night), recycling nitrate into ammonium when aerobic nitrification is less active. This ensures stable ammonium availability, potentially supporting coral growth. In contrast, coupling of nitrification and denitrification may help regulate nitrogen levels under nutrient-rich conditions, preventing excessive proliferation of Symbiodiniaceae and maintaining a balanced holobiont metabolism.

Overall, our results expand current understanding of nitrogen cycling in coral holobionts by resolving the microbial taxa actively involved and mapping their compartment-specific activities. The coral skeleton emerges as a metabolically active zone for both aerobic and anaerobic nitrogen transformations, with stratified microbial communities performing complementary functions. These findings underscore the importance of skeleton-associated microbes in holobiont nutrient dynamics and reef ecosystem functioning.

### Sulphur metabolising microbes are widespread across coral compartments

Sulfur, abundant in seawater primarily as inorganic sulfate, is assimilated by the coral host, Symbiodiniaceae, and various members of the coral microbiome into essential organic compounds, making it a critical element for holobiont growth and function (*92–94*). In this study, we identified expression of transcripts encoding sulfur cycling marker genes across all three coral compartments (Fig. 3B), underscoring the distributed nature of sulfur metabolism within the holobiont. Notably, transcript abundance for most sulfur-related genes was highest in the skeleton, indicating a prominent role for the endolithic microbiome in coral sulfur cycling.

We detected expression of anaerobic dissimilatory sulfate reduction (DSR) marker genes, aprA and aprB, exclusively in the skeleton compartment. Taxonomic annotation identified these transcripts as originating from canonical sulfate-reducing bacteria (SRB), including Deltaproteobacteria, as well as anoxygenic phototrophic Chlorobi (Fig. 3B). Although Chlorobi carry the aprAB genes, they typically participate in sulfur oxidation rather than sulfate reduction (95, 96), and their presence here suggests diverse sulfur redox processes within the skeleton. This observation supports prior metabarcoding and metagenomic studies, which reported abundant SRBs and assembled their genomes from the skeleton of *P. lutea* and *I. palifera* (24). The co-detection of SRBs and Chlorobi within the same compartment aligns with previous reports of anoxic microniches in coral skeletons (*14*, *24*, *27*, *76*) and suggests the presence of syntrophic interactions, wherein SRB-produced sulfide serves as an electron donor for phototrophic Chlorobi. Such metabolic coupling has been observed in microbial mats, stratified lake waters, and coral skeletons (*27*, *97–99*). The simultaneous expression of aprAB genes by both groups in this study provides molecular evidence for this syntrophy within the coral holobiont.

We also identified a complex and spatially distributed expression pattern of **assimilatory** sulfate reduction (ASR) genes. Transcripts for cysC, cysI, cysJ, and SUOX were detected across all compartments, whereas cysD expression was restricted to the deeper skeleton (Fig. 3B). The gene encoding cysK, which links assimilatory sulfate reduction to L-cysteine biosynthesis, was consistently expressed in all compartments. These findings suggest widespread ASR activity throughout the holobiont, supporting sulfur assimilation and amino acid biosynthesis. Taxonomic analysis of ASR gene transcripts revealed a diverse array of eukaryotic and prokaryotic symbionts. These included microeukaryotes such as Symbiodiniaceae, *Ostreobium*, and other Chlorophyta, as well as bacteria including Cyanobacteria, Thaumarchaeota, and Proteobacteria (Fig. 3B). Notably, all ASR transcripts were exclusively assigned to algae and bacteria, consistent with the known distribution of this pathway among plants, algae, and microbes(*100*).

Together, these results provide new insights into the compartmentalized and microbially mediated sulfur cycling processes within *P. lutea*. The coral skeleton emerges as a key site for both dissimilatory and assimilatory sulfur transformations, driven by specialized microbial consortia. The detection of potential syntrophic sulfur interactions between SRBs and Chlorobi adds a new layer of complexity to our understanding of the coral skeleton as a metabolically active niche, reinforcing its contribution to holobiont nutrient cycling and resilience.

### Microeukaryotes are the major scavengers of ROS and RNS in *P. lutea*

Coral reefs are facing unprecedented threats due to rising sea surface temperatures, resulting in more frequent and prolonged coral bleaching events (*37*). Most theories on the onset of coral bleaching suggest that the overproduction of reactive oxygen species (ROS) and reactive nitrogen species (RNS) is a key trigger, leading to cellular damage and stress under adverse environmental conditions (*101*). Rapid scavenging of ROS and RNS is thus crucial in maintaining homeostasis and protecting the coral holobiont in stressed and healthy corals. We observed widespread expression of seven protein families with functions related to ROS or RNS scavenging across the tissue, *Ostreobium* layer, and deeper skeleton compartments. Specifically, transcripts encoding for genes involved in RNS scavenging, including nitric oxide reductases (PF00175: *hmp*; PF00115 and PF00034: *nor*BC) and peroxynitrite to nitrate reduction (PF10417: *ahp*C) were expressed in all compartments (**Fig 4**). A single transcript encoding nitrous oxide reductase (PF18764: *nos*Z) was identified and expressed exclusively in deeper skeleton samples. This enzyme catalyses the last step of the dissimilatory metabolic pathway of denitrifying bacteria, which were abundant in the deeper skeleton. Similar observations were made for ROS scavenging expression profiles. We identified 48, 43, and 41 transcripts encoding the general ROS scavenger Glutaredoxin (PF00462), as well as hydrogen peroxide scavengers Peroxiredoxins (PF00578) and peroxidases (PF00141). These transcripts were expressed in all coral compartments along with transcripts encoding superoxide dismutases (PF00081 and PF02777: SOD Mn and PF00080:SOD cu-zn) (**Fig 4**). The majority of ROS and RNS scavenging transcripts were of micro-eukaryotic origin (76%), including Symbiodiniaceae, *Ostreobium*, Chlorophyta, and Fungi, while others were from archaeal and bacterial sources (24%), encompassing Thaumarchaeota, Proteobacteria (Rhodospirillales, Xanthomonadales), Cyanobacteria, Chlorobi, Chloroflexi and Bacteroidetes (*Candidatus* Amoebophilus) (**Supplementary data file 4**).

**Fig. 4.**
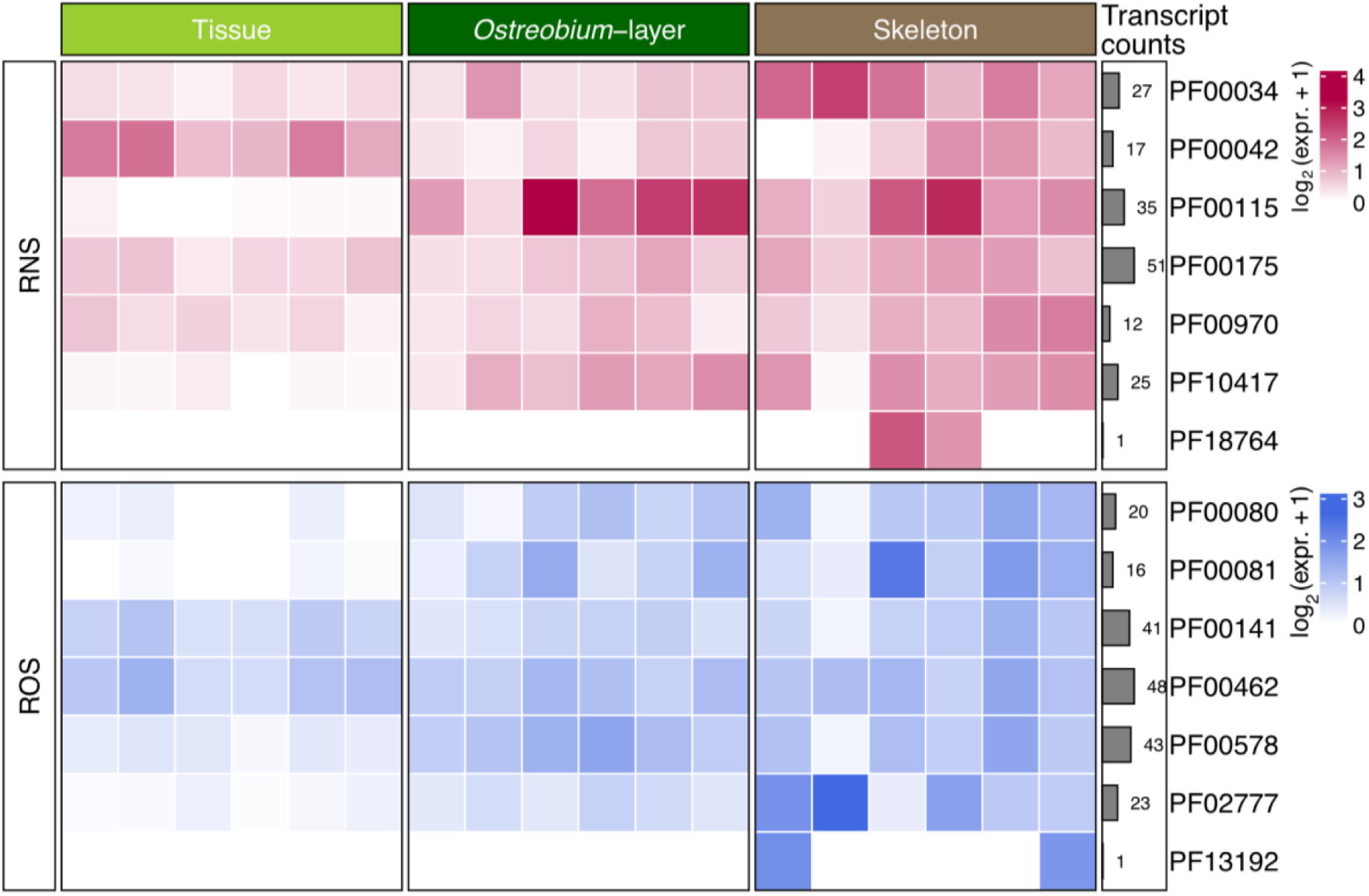
Transcriptome-based expression profiles of Reactive Oxygen Species (ROS) and Reactive Nitrogen Species (RNS) marker genes across coral compartments. Heatmap displaying the expression of different protein families, count of transcripts for each family involved in ROS and RNS scavenging across coral compartments. Expression values for both were averaged and log2(expression + 1) transformed.

Scavenging reactive oxygen (ROS) and nitrogen species (RNS) is a key function of bacterial candidates for coral probiotics, as emphasised in recent studies identifying potential probiotic bacteria for corals (*2*, *26*, *102–104*). However, the identification of the majority of ROS and RNS scavenging transcripts in micro-eukaryotic inhabitants of the coral holobiont rather than in bacteria, raises the question of which organisms are the main players in scavenging ROS and RNS under thermal stress. This finding suggests that current strategies for designing coral probiotics, which primarily focus on bacterial functions, should be refined through additional molecular and experimental validation to provide functional insights. These insights could guide the assembly of a more comprehensive consortium of beneficial symbionts for corals.

## Conclusion

This study provides novel insights into the transcriptionally active communities within the coral holobiont, with a focus on their functional roles across distinct ecological compartments: coral tissue, the endolithic *Ostreobium* layer, and the deeper skeleton layer. Our analysis revealed a nuanced stratification of microbial activity, with microeukaryotes dominating key metabolic processes in the coral tissue and *Ostreobium* layer, while prokaryotes emerge as the primary active symbionts in the deeper skeleton (**Fig 5**). These findings underscore the profound influence of physicochemical gradients—including light penetration, nutrient availability, and carbon distribution—in sculpting both community composition and functional specialization within the holobiont. Functional analysis highlighted the critical contributions of *Symbiodiniaceae, Ostreobium*, and Cyanobacteria to photosynthate production, with *Symbiodiniaceae* predominantly occupying the coral tissue, while *Ostreobium* and Cyanobacteria play pivotal roles in the endolithic *Ostreobium-*layer (**Fig 5**). These symbionts, evolutionarily adapted to varying light conditions and utilizing distinct photosynthetic pigments, demonstrated unique nutrient cycling capabilities. Notably, in addition to *Symbiodiniaceae*, we identified *Ostreobium* and fungi as the only taxa capable of exporting sugars via SWEET proteins, providing novel insights into nutrient exchange mechanisms within the holobiont. Analysis of nitrogen metabolic pathways revealed Cyanobacteria as the exclusive nitrogen fixers, with other nitrogen and sulphur metabolic processes likely distributed across holobiont members, suggesting a robust functional redundancy that potentially enhances ecosystem resilience (**Fig 5**). Our analysis also identified microeukaryotes as the primary scavengers of ROS and RNS across all compartments, playing a crucial role in maintaining holobiont health, particularly under thermal stress conditions (**Fig 5**). These findings offer a comprehensive view of the metabolic activities within the coral microbiome, elucidating the intricate functional roles of its diverse members. While this research significantly advances our understanding of coral holobiont functions, it simultaneously highlights the complexity of these systems and the necessity for more targeted research focussed on the microenvironmental niches of the coral holobiont. Future investigations should employ advanced sampling techniques and high-resolution analytical methods to more precisely map the functional dynamics between coral hosts, their symbionts, and the surrounding microenvironment (*11*, *105*, *106*). Ultimately, the insights gained from this study are invaluable for advancing coral reef conservation efforts. By identifying functionally relevant symbiotic members, we can better inform the design of coral probiotics and other strategies aimed at enhancing coral resilience in the face of climate change and other environmental stressors.

**Figure 5.**
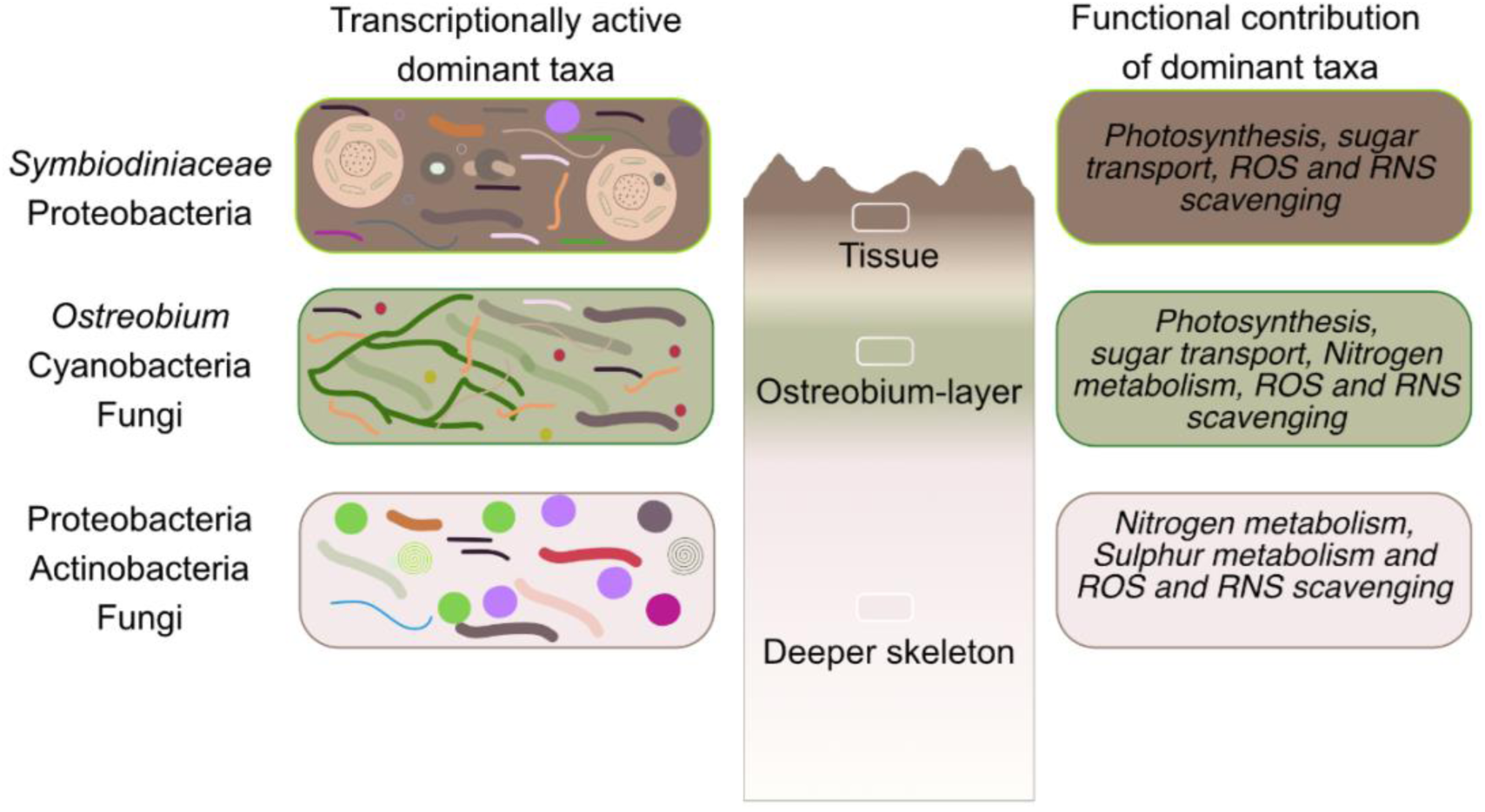
Overview of the transcriptionally dominant taxa and their functional contribution in different compartments of the coral identified in this study.

## Data and code availability

Raw reads generated in this study have been submitted to NCBI SRA under the Bioproject PRJNA1207708. All codes and files required to regenerate the figures have been made available via github.

## Acknowledgement

Authors would like to acknowledge staff at the Heron Island Research Station for their assistance during sample collection. All coral samples were collected under Permit G22/47504.1 issued to KT and FR by Great Barrier Reef Marine Park Authority, Australia. Authors would also like to acknowledge reviewers of the manuscript for their suggestions.

